# Chemical probes to interrogate the extreme environment of mosquito larval guts

**DOI:** 10.1101/2023.12.27.573438

**Authors:** Lindsay E. Guzmán, Anjalee N. Wijetunge, Brendan F. Riske, Brooke B. Massani, Michael A. Riehle, John C. Jewett

**Affiliations:** Stanford University; University of Arizona

## Abstract

Mosquito control methods are vital for the spread of life-threatening illnesses such as dengue fever, malaria, and yellow fever. Vector control technologies must be selective to minimize deleterious effects to our ecosystem. Successful methods that control mosquito larva populations utilize the uniquely high alkaline nature of the midgut. Here, we present novel protected triazabutadienes (pTBD) which are deprotected under basic conditions of the larval midgut, releasing an aryl diazonium ion (ADI) that results in protein modification. The probes contain a bioorthogonal terminal alkyne handle, enabling a selective Cu-click reaction with an azido-fluorophore for quantification by SDS PAGE and visualization using fluorescence microscopy. A control TBD, unable to release an ADI, did not label the midgut. We envision our chemical probes will aid in the development of new selective mosquito control methods thus preventing the spread of mosquito-borne illnesses with minimal impact on other organisms in the ecosystem.

## Introduction

Mosquito-borne pathogens are responsible for over 500,000 deaths annually with malaria parasites and arboviruses (from anopheline and culicine mosquitoes, respectively) being primary culprits.^1^ Current mitigation strategies rely heavily on vector control of these mosquito subfamilies at two key life stages, the larvae and adult (**Fig. 1A**). However, commonly used mosquito adulticides and larvicides are becoming ineffective as resistance develops.^2^ For current chemical and biological mosquito larvicides *only four* modes of action (MOA) are widely utilized, acetylcholinesterase inhibitors (temephos), nicotinic acetylcholine receptors (spinosad), juvenile hormone mimics (methoprene and pyriproxyfen) and microbial disruptors of insect midgut membranes (*Bacillus thuringiensis israelensis* (BTi) and *Bacillus sphaericus* (BS)).^2a, 3^ Of these, resistance has been reported in various mosquito species against temephos, spinosad and pyriproxyfen.^4^ Field resistance against BTi has been low, likely due to BTi formulations containing four different toxins, greatly mitigating the likelihood of resistance developing. In contrast, BS produces a single toxin and multiple instances of resistance have been reported worldwide.^5^ Thus, there is a critical need for novel larvicides leveraging additional MOAs and companion tools to probe larval mosquito physiology.

**Figure 1.**
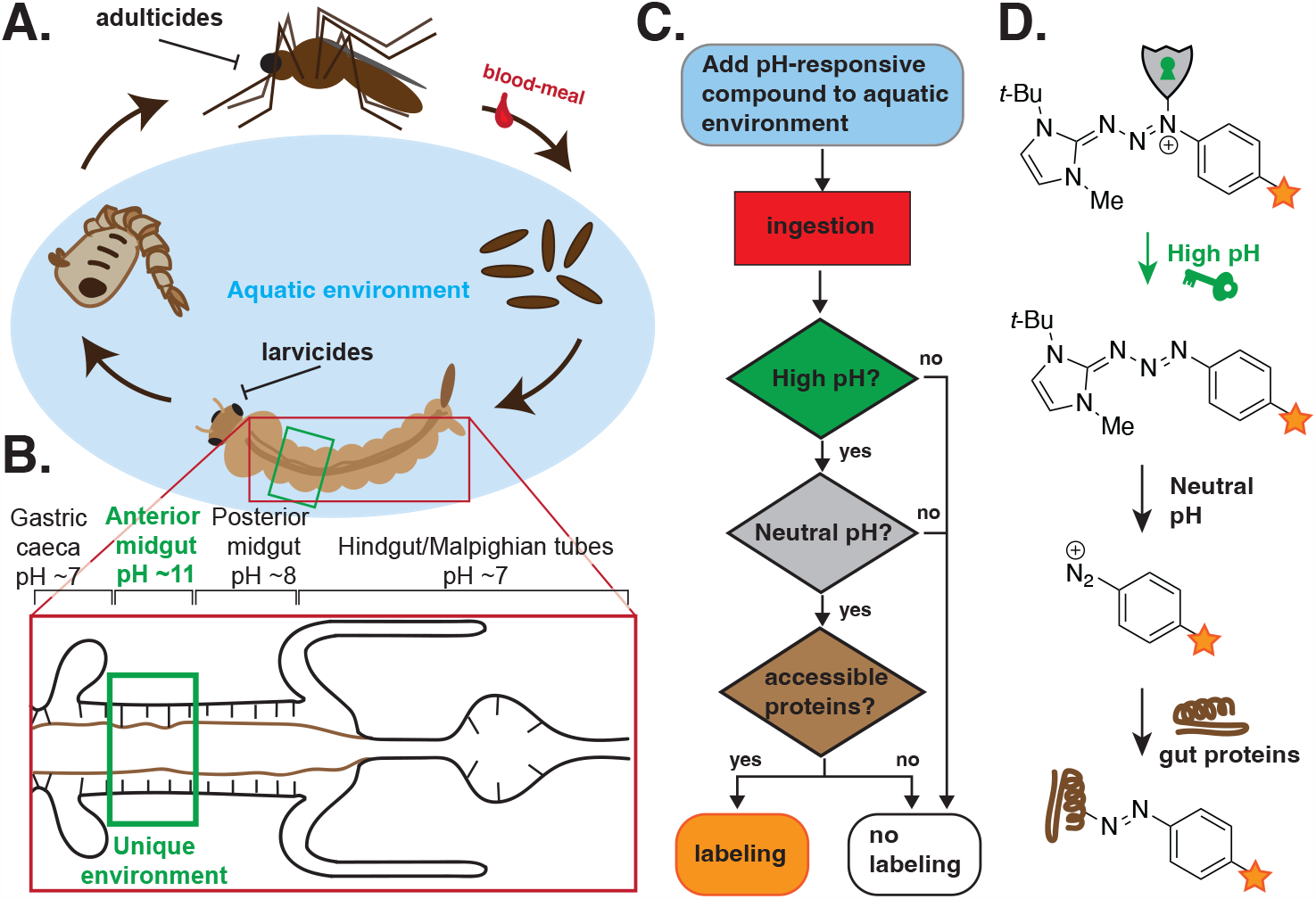
**A**) Mosquito life-cycle. After taking a blood-meal adult female mosquitoes deposit eggs in water or a suitable oviposition substrate. Following embryo development the larvae emerge from the egg, develop through four larval instars, pupate, and eclose as adults. **B**) The alimentary tract of the mosquito larva is highlighted to show the uniquely basic environment of the anterior midgut. **C**) We seek a molecule that will respond to the variable pH midgut environment and only label midgut proteins when the “yes” conditions being met. **D**) Triazabutadienes with base-labile protecting groups serve to release aryl diazonium ions which can covalently modify proteins.

An ideal mosquito larvicide should be adaptable to changes in resistance and highly specific to minimize non-target effects. The specificity of BTi is due in part to it only becoming soluble and active in the unique alkaline environment of the larval mosquito midgut. Only a handful of organisms, including immature stages of some dipteran (flies), coleopteran (beetles) and lepidopteran (butterflies and moths) species, provide such an alkaline environment.^6^ Thus, leveraging this unique biology ensures a highly specific larvicide with minimal off-target effects, particularly in the aquatic environment of the larval mosquito. The key target for ingested larvicides is the mosquito alimentary system, and specifically its midgut where digestion and nutrient absorption occurs. The larval mosquito midgut can be further divided into three sections, the sac-like gastric caeca,^7^ anterior midgut, and posterior midgut (**Fig. 1B**). The anterior larval midgut, a region responsible for the initial stages of protein and lipid breakdown and lipid absorption, maintains an exceptionally high pH of 10.5-11.5 in various mosquito species, including the important vector species *Ae. aegypti* and *A. stephensi*.^8^ The posterior midgut gradually decreases in pH to a final value of ∽8 prior to excretion.

Lining the food bolus is the peritrophic matrix (PM), which protects the midgut and establishes a counter-current flow back to the GC for additional nutritional absorption.^9^ Inspired by this unique chemical environment, we designed molecules capable of targeting and modifying larval midgut proteins when exposed to the variable pH environment of the larval gut, schematized in **Figure 1C**, the first step of a long-term goal to deliver novel larvicidal agents. In this study we report a class of molecules that require a *sequential* two-stage activation prior to releasing an electrophilic aryl diazonium ion that covalently modifies larval gut proteins (**Fig. 1D**).

Prior to realizing the goal of larvicides with novel MOAs, we sought to develop and establish a chemical approach that navigates the chemistry of the larval gut and covalently modify proteins in the midgut lumen. Based on our previous work with protected triazabutadienes,^10, 11^ and their delivery of protein-reactive aryl diazonium ions (ADIs),^12^ we hypothesized that if mosquito larvae were exposed to these compounds, the probes would remain protected until reaching the basic anterior midgut, which would initiate the release a reactive ADI to modify surface-exposed proteins (**Fig. 1D**). To test that hypothesis, the chemical probes (**1, 2** and control **3, Fig. 2A**) were designed to take advantage of the variable pH environment of the larval gut, enabling detection of midgut proteins and disruption of essential midgut physiologies. In evaluating the compound design logic from ingestion to detection (**Fig. 1C&D**), the first requirement is stability in the aqueous environment in which larvae develop. When the nucleophilic N3 nitrogen (**Fig. 2B**) of the triazabutadiene reacts with certain electrophiles it prevents them from protonating at the N1 nitrogen, a requisite for ADI liberation.^13^ Two different carbamates were chosen, both expected to hydrolyze in the basic environment, but varied in structure. An alkylated version of the probe serves as a structurally similar control and should not release the triazabutadiene, or subsequent diazonium ion due to its stability to both acidic and basic environments.^13^ Triazabutadienes derived from a *t*-butyl methyl imidazolium ion were previously shown to release rapidly at neutral pH,^14^ and have proven useful with both labeling of proteins^15^ and delivery.^11^ Finally, we installed an alkyne for downstream detection on all of the chemical probes. The modular nature of a terminal alkyne provides a bioorthogonal Cu-click handle that can facilitate linkage to a wide range of azide-containing molecules for future studies.^16^ Terminal alkynes are exceptionally rare in nature, and we hypothesized that larval guts would not have cross-reactivity with azidofluorophores.

**Figure 2.**
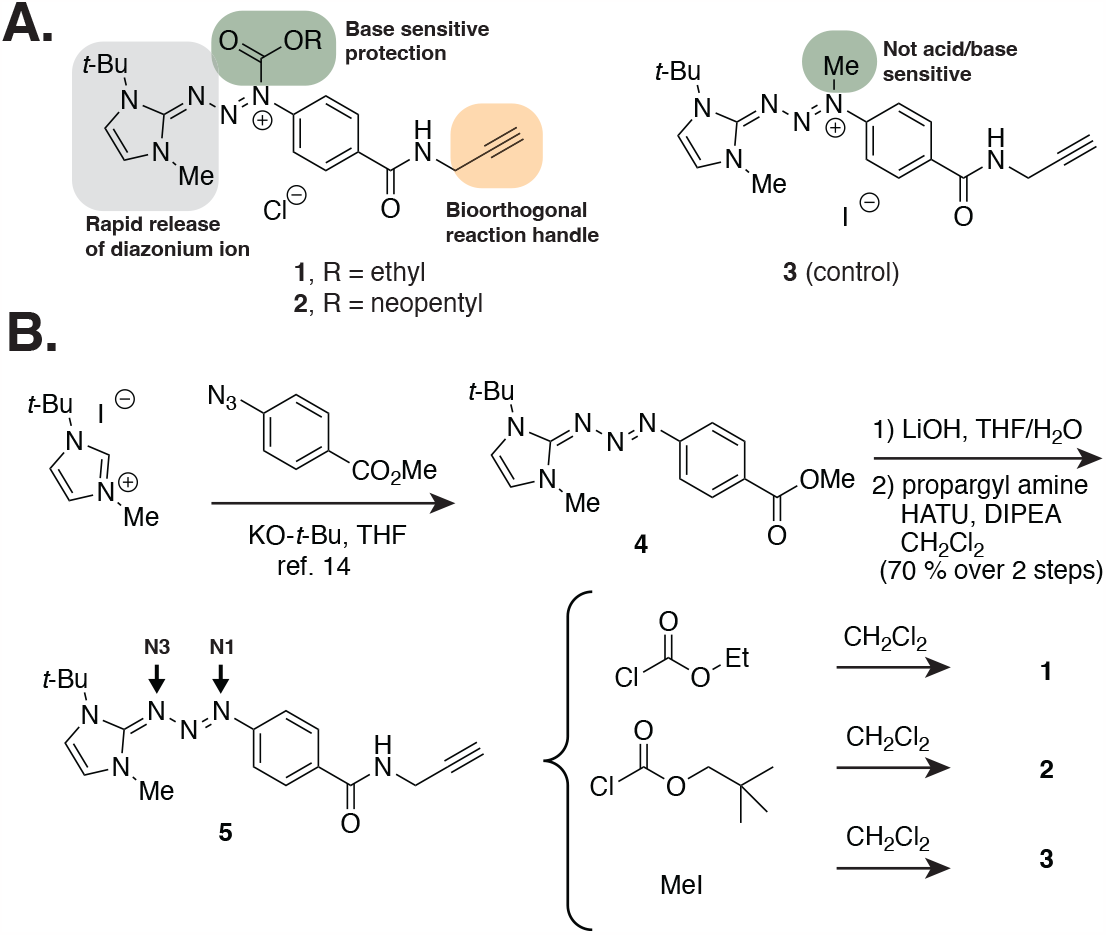
**A)** Design features for mosquito larva probes. **B**) Synthesis of experimental and control molecules.

### Discussion and Experimental Section

Following established synthetic protocols,^15,17^ the triazabutadiene core, **4**, was made using an in-situ generated carbene and aryl azide, synthesized from 4-aminobenzoic acid (**Fig. 2B**). Hydrolysis of the methyl ester and coupling to propargyl amine provided triazabutadiene **5** in good yield from ester **4**. Protection of the triazabutadiene was accomplished with ethyl chloroformate, neopentyl chloroformate, or methyl iodide to provide base-reactive compounds **1** and **2**, or control compound **3**, respectively. It was previously established that ethyl and neopentyl carbamate protected triazabutadienes provide similar release kinetics.^10^ Indeed, small molecule kinetics experiments showed complete consumption of **1** and **2** over less than 10 minutes at pH 10, while at pH 7 no changes were observed over an hour. Compound **3** remained unchanged in either environment (see Supporting Information **Fig. S1**).

We chose to work with the mosquito *Aedes aegpyti*, a widespread and important disease vector capable of transmitting a range of arboviruses that impact human health, including dengue, Zika, yellow fever and Chikungunya viruses. Mosquito larva transition through four development stages (called instars) over the course of ∽8 days, growing larger in each stage. Once the larvae have acquired sufficient nutrients they undergo metamorphosis, resulting in a dramatic reorganization of proteins and tissues during the pupal stage.^18^ We initiated to start our experiments with late 2^nd^-early 3^rd^ instar larvae and analyzed their midguts during the early to mid 4^th^ instar. This provided sufficient time for the larvae to ingest the compounds via filter feeding, and allowed us to work with the largest instar to facilitate midgut dissection and probe detection. Most protocols for protein isolation are optimized for tissue samples and proteins that are adapted to neutral or acidic environments, and we found that standard protease inhibitor cocktails (such as Halt^TM^ protease inhibitor) failed to sufficiently inhibit protein degradation as assessed by the presence of degraded, low molecular weight proteins using SDS-PAGE analysis (**Fig. 3A** and see Supporting Information, **Fig. S2**).^19^ Other studies also reported a low yield of identified proteins from *Ae. aegypti* midguts dissected in neutral buffer.^20^ Coupled with our observation, we inferred that proteolytic degradation during dissection due to the abundance of digestive enzymes was the likely culprit. Indeed, transcriptomic analyses revealed that *Ae. aegypti* have 66 genes encoding “trypsin-like” serine proteases, which are principally expressed and highly active in the larval midgut.^21^ Studies showed that larval digestive enzymes are inhibited in a highly acidic buffer solution,^22^ but these insights had not been previously applied to lysates/dissections. Therefore, we dissected mosquito larvae in the presence of 0.1 M sodium acetate at pH 3 and paid careful attention to pH throughout tissue processing. Using this approach we were able to isolate intact proteins that resolved well on SDS PAGE (e.g., **Fig. 3C**).

**Figure 3.**
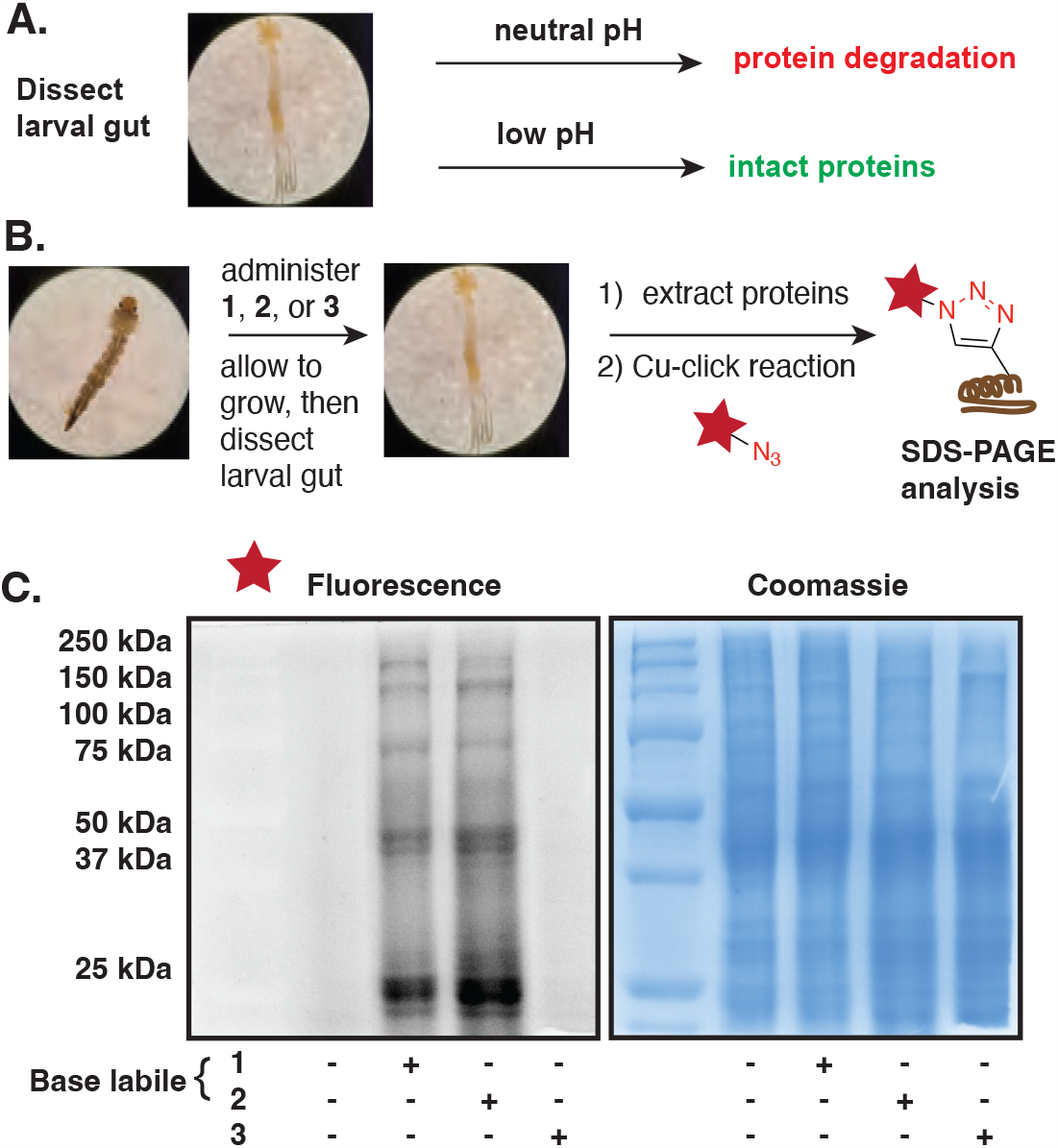
**A**) Larva must be dissected in acidic media to deactivate proteolytic enzymes. **B**) Mosquito larva are grown in water containing compounds **1, 2**, or **3** for 16 hours in an insectary. The 4^th^ instar larvae were dissected and their guts were either used for protein detection or imaging experiments. Covalently modified proteins were detected by Cu-click with fluorescently labeled azides. **C**) Larva that had grown in the presence of **1** and **2** showed a range of proteins had been modified. Control compound **3** did not modify proteins.

The general experimental workflow is highlighted in **Figure 3B**. To assess probe-dependent labeling of the mosquito larvae, cohorts of 30-40 2^nd^ or 3^rd^ instar *Ae. aegypti* larvae were incubated with 1 mM solutions of compounds **1, 2** or **3** (control) in tap water for 16 hours.^23^ As discussed above, larva were dissected in acidic buffer to minimize protease activity and dissected midguts were snap frozen in liquid nitrogen prior to protein extraction by homogenization and dissolution in lysis buffer.^24^ Midgut lysates were subjected to Cu-click reaction conditions in the presence of AlexaFluor^TM^ 555 azide and then analyzed by running on SDS-PAGE (**Fig. 3C**). As envisaged, significant fluorescent labeling was observed in the protein lysates from mosquitoes treated with either of the experimental compounds (see Supporting Information **Fig. S3** for densitometry analysis, Coomassie staining was used to verify equal protein loading in all lanes). In contrast minimal, non-specific fluorescence was observed in untreated mosquitoes or those treated with the control compound. This demonstrates the expected workflow (shown in **Fig. 1D**) consisting of multistep deprotection followed by bioconjugation, was accomplished in the larval gut during larval filter feeding. In accordance with a basic hydrolysis mechanism to provide the same triazabutadiene and resulting ADI, the labeling for both ethyl and neopentyl protected compounds was robust (**Fig. 3A**). Moreover, the lack of statistically significant labeling with the alkylated triazabutadiene, **3**, as compared with a non-treated sample minimized the likelihood that non-specific binding was a culprit for labeling. Interestingly, neopentyl protected **2** was shown to have a small, but statistically significant, increase in labeling as compared with ethyl protected **1** (see Supporting Information **Fig. S3**). This difference may result from differing pharmacokinetic properties of the intact probes, and will be the subject of future investigations. Notably, none of these probes resulted in noticeable larval toxicity at the concentrations and times tested, demonstrating that protein modification with these probes alone is not acutely toxic during this exposure time. That said, some fitness effects (e.g., increased development time, reduced growth, reduced pupation rate, etc.) may occur that we did not detect in our initial assays may occur with increased exposure to the reactive compounds. Accordingly, future studies will explore the impact of various exposures for a variety of mosquito life history traits. Nevertheless, this provides an exciting opportunity to leverage a range of mosquitocidal-specific payloads as a future mechanism of toxicity and larval control.

Once we established that these compounds label proteins in the larval midgut we turned attention to imaging how pervasively the gut was labeled (**Fig. 4A&B**). Similar to the lysate studies above, larvae were incubated with 1 mM solutions of compounds **1, 2**, or **3** (control) overnight in a controlled insectary environment. Larval midguts were dissected in low pH buffer, the guts were washed extensively to remove unreacted compounds, fixed with paraformaldehyde and permeabilized with Triton-X detergent.

**Figure 4.**
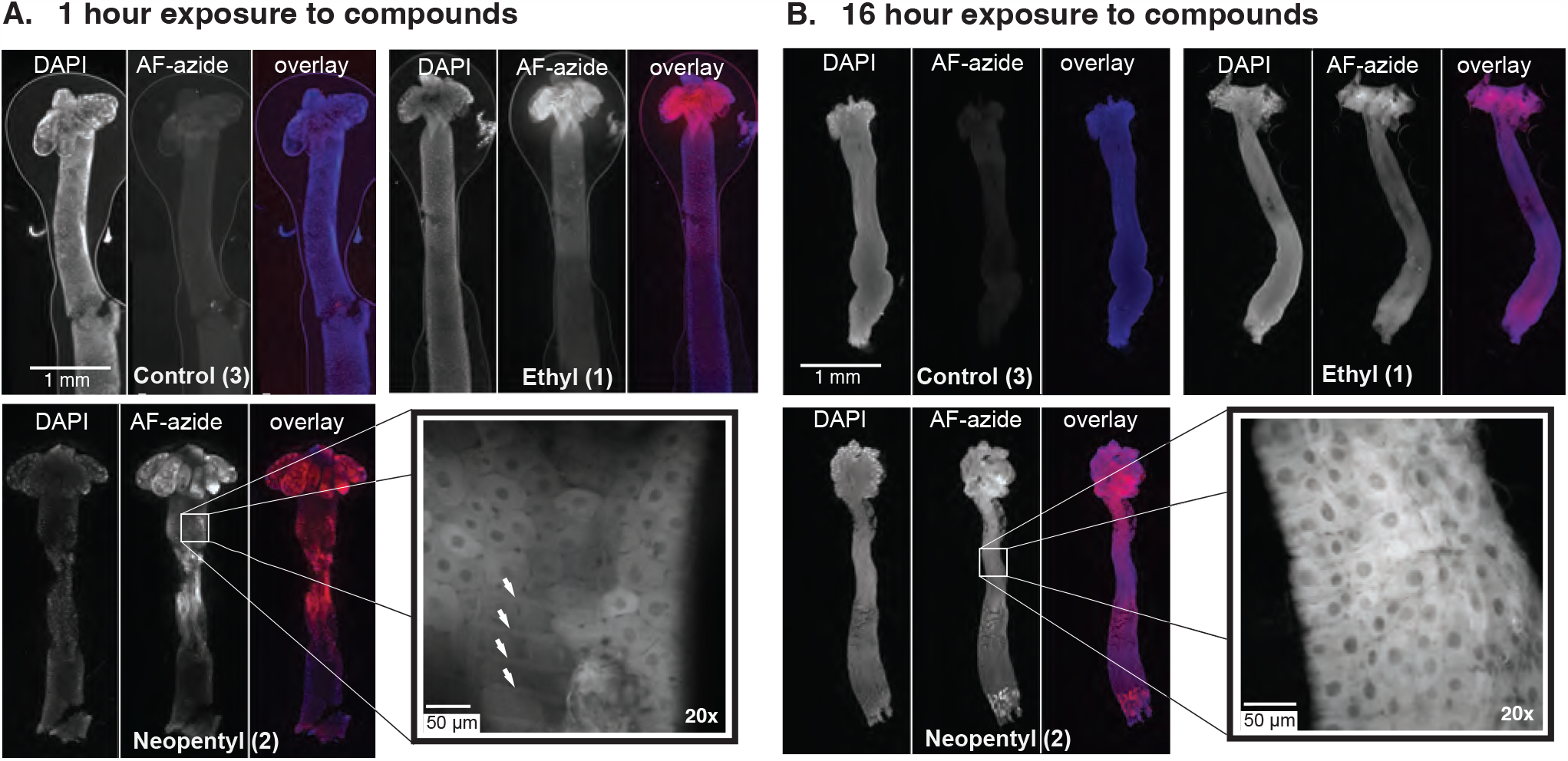
After following the same treatment/dissection workflow from **Fig. 3A**, larval guts were fixed, permeabilized and subjected to Cu-click conditions with an AF-555 azide plus dye. The whole body images are 5x, obtained with an epifluorescence filter, and confocal microscopy was used to obtain the 20x inset images. The overlay images show DAPI in blue, and AF-azide in red. **A**) Labeling occurred after only 1 hour of exposure, but was more variable. Muscle fibers can be seen, highlighted by white arrows. **B**) Robust labeling of the entire alimentary tract of larval guts with compounds **1** and **2** was observed by fluorescence microscopy after 16 hours of exposure.

The midguts were treated with Cu-click conditions using an AlexaFluor^TM^ 555 azide plus^25^ fluorophore and then imaged using epifluorescence and spinning-disk confocal microscopy. The imaging results (**Fig. 4B**) were consistent with what was observed with the SDS-PAGE analysis of protein lysates with strong fluorescence observed in mosquitoes treated with experimental compounds (**1 & 2**) and minimal fluorescence observed with either the control compound (**3**) or the non-treated samples. This negates the likelihood of non-specific fluorophore labeling. Interestingly, the entire alimentary tract was labeled, likely due to the nutrient flow in larval guts where small molecules/nutrients can flow in a counter current opposite the food bolus to eventually reach the gastric caeca for increased adsorption opportunities. Our small molecule kinetics experiments hinted that we *should* be able to observe labeling at an earlier timepoint, and thus followed the workflow above with larva exposed to compounds **1**-**3** for just 1 hour (**Fig. 4A**). The resulting images showed unevenly labeled regions, with sample to sample variability (see Supporting Information **Fig. S4**), likely due to the inherent variability in each larvae that was mitigated over the longer time course.

Compound **2** generally showed better labeling at 1 hour, again hinting at a pharmacokinetic difference. Interestingly, different tissues, such as muscle fibers^26^, were clearly labeled at 1 h, but not at the later timepoint. This could be an artifact of muscular fiber remodeling that occurs, or simply the other cells were significantly brighter due to more labeling and the faint muscle fiber labeling was not visible. While the luminal side of the midgut did appear to be labeled more than the basal side exposed to the hemolymph (see **Supporting Videos 1-2**), as might be expected in this system, the labeling of muscle fiber clearly shows that ADI probes are capable of passing through the larval gut cells and targeting proteins on the basal side. This finding could prove beneficial if potentially toxic compounds need to be delivered intracellularly or to the hemolymph. The widespread labeling offers intriguing possibilities for larvicides that leverage a base-promoted pro-larvicidal strategy. Future studies examining larvicidal activity and time-course or trafficking of labeling are planned but will require a new generation of probes. Additionally, work is currently underway to identify chemical probe targets by proteomics.

## Conclusions

We reported a chemical approach to covalently modify gut proteins of *Ae. aegypti* mosquito larvae leveraging their unique high pH environment as a triggering mechanism. We describe our design, synthesis, and application of our chemical probes. We demonstrated the efficacy of our probes using SDS PAGE and fluorescence microscopy. The procedures and methods developed will be useful to entomologists as an additional proteomic analysis tool during various larval stages that may help identify candidate drug targets to develop new insect control programs. In the future, we envision that our technology will aid in the development of new mosquito larvicides with a high degree of target specificity and the ability to rapidly adapt to changes in field resistance, and thus limit the spread of mosquito-borne illness. Finally, the ability to systematically imbue a molecule with pre-programmed logic to match a biological environment should prove broadly useful beyond mosquito larva and their uniquely basic gut.

## Acknowledgements

The authors would like to thank Dr. Garrett Davis for biochemical discussions, Dr. Natasha Cornejo for synthetic discussions, and Jenet Soto-Shoumaker for larval maintenance. The material is based upon work supported by the National Science Foundation under Grant No. CHE 1552568 to JCJ, supporting synthesis and early biochemical studies. Support for larval studies was provided by the 2023 Technology Research Initiative Fund (TRIF)/ BIO5/Improving Health Seed grant administered by the University of Arizona Office for Research, Innovation and Impact (A.R.S.§15-1648). The imaging studies were carried out with funding support from NIH grant R21AI177824 to JCJ & MAR. LEG thanks the Maria Teresa Velez Diversity Fellowship Award 2017-2018, and NIH Training Grant (T32 GM008804). All fluorescence images and data were collected in the W.M. Keck Center for Nano-Scale Imaging in the Department of Chemistry and Biochemistry at the University of Arizona, RRID:SCR_022884. This instrument purchase was partially supported by Arizona Technology and Research Initiative Fund (A.R.S.§15-1648).

