## Supplemental information for "Chemical probes to interrogate the extreme environment of mosquito larval guts"

University of Arizona, Tucson, AZ 85721

### Supporting information

### General information

The chemicals including 4-aminobenzoic acid, concentrated HCl, sodium nitrite, sodium azide, potassium *tert*-butoxide, 1-*tert*-butyl imidazole, methyl iodide, LiOH, diisopropylethylamine, *N*-hydroxysuccinamide, HATU, triethyl amine, propargyl amine, chloroformates, $CuSO_4 \cdot 5H_2O$, $CuCl_2$, sodium ascorbate, THPTA and all the solvents and chromatography materials including silica, neutral alumina and basic alumina were purchased commercially and used as received.

NMR spectra were taken on a Bruker AVIII-400 NMR Spectrometer, Bruker AVIII-500 NMR or Avance NEO 500 for $^1H$ and $^{13}C$ NMR and referenced with residual solvent peaks at 7.26 ppm and 77.0 ppm for $CDCl_3$, 2.50 ppm and 39.5 ppm for DMSO-$d_6$. Coupling on the NMR spectra is expressed in Hertz (Hz), chemical shifts in ppm, with abbreviations for multiplicities as s = singlet, d = doublet, t = triplet, q = quartet, p = pentet, m = multiplet. Mass spectra were obtained using solariX 2XR 9.4T FTICR instrument. Mass spectral analysis was performed on a Bruker ICR ESI. Infrared spectra were obtained using a Thermo Fisher Scientific Nicolet IS50R FT-IR Spectrometer. The transmission mode was attenuated total reflectance (ATR), using a Spectra-Tech Thunderdome germanium crystal ATR accessory. The detector was a liquid nitrogen cooled high sensitivity MCTA detector. The FT-IR signals are reported as w = weak, m = medium s = strong, br = broad. The single crystal XRD measurements were performed at the XRD facility in the Department of Chemistry and Biochemistry of the University of Arizona, on a Bruker Kappa Apex II X-ray diffractometer. The molybdenum Ka X-ray source was used, and the measurement temperature was 100 K. Absorbance spectra were obtained using an Agilent 8453 UV-visible spectroscopy system. Epifluorescence and spinning disk confocal images were taken using an Intelligent Innovations Imaging (3i, Denver Colorado) configured Zeiss Marinas microscope in the W.M. Keck Center for Nano-Scale Imaging, RRID:SCR_022884 (Figure 4). Images of the whole dissected mosquito larval gut were collected using a 5X EC Plan-Neofluar objective, DAPI filter, CY3 filter, and Andor DU 897 EMCCD. A one by three or one by five montage image array was collected. The images were stitched together using SlideBook 6.0.23 software to enable visualization of the whole larval gut. For each experiment the same collection settings were used for each sample (ethyl (**1**), neopentyl (**2**), control (**3**), and non-treated larval guts). Images comparing the different samples for each experiment were adjusted such that the LUT for each image was the same using neopentyl (**2**) sample to define the maximum intensity. Spinning disk confocal fluorescence images were collected using the same microscope base with the spinning disk confocal optical path. Images were collected using a 20X EC-Plan-Neofluar objective, Yokogawa CSU-X1 M1 with dual-emitter dichroic and bandpass filters centered around 525/43 nm and 641.5/117 nm, and a Photometrics Evolve 512 EMCCD. Movies were made using spinning disk confocal Z-stack images collected using 0.630 µm step size and exported using SlideBook 6.0.23 software with 0.2 sec frame duration.

BSA stocks were made from commercially purchased lyophilized powder from Sigma Aldrich (9048- 46- 8). SDS-gels were conducted with freshly poured 10 % SDS gels. Precision Plus Protein™ All Blue Prestained Protein Standards (Bio-Rad : 1610373) was used as the molecular weight marker during gel electrophoresis. 6X Laemmli sample buffer (1.2 g SDS, 4.7 mL glycerol, 6 mg bromophenol blue, 1.2 mL 0.5M pH 6.8 Tris, 2.1 mL distilled water and 0.93 g DTT) was used in the respective amounts as mentioned in the protocols. AZDye™ 555 Azide (Click Chemistry Tools : 1287-1) and AFDye 555 Azide plus (Click Chemistry Tools) were used in the in-gel fluorescence and imaging experiments. Click reactions with fluorophores and subsequent steps before imaging were performed void of light. Fluorescent gel imaging was performed with a ChemiDoc™ MP scanner (BioRad : Pharos FX Plus Molecular Imager). Respective laser lines and the filters were used to image AZDye™ 555 Azide : 546 nm UV-laser and 605/50 nm filter. Coomassie staining was imaged using the same scanner. Fluorescence images were obtained using the Spinning Disk Confocal/TIRF/Widefield microscopes from Intelligent Imaging Innovations in the CBC Keck facility and processed using the slide book software. ProLong™ Gold Antifade Mountant with DAPI for larval gut tissue mounting was purchased from Thermo Fischer Scientific (P36941). Mosquito larval dissections were performed using Nikon SMZ-10A microscope.

Mosquito Rearing*:*

*Aedes aegypti* mosquitoes were maintained in an insectary at 27 °C and 75% relative humidity in a 16/8 h light/dark photoperiod. Larvae were reared at a density of 150 individuals per liter of water and fed ground cat chow (Purina Cat Chow Complete). Pupae were transferred into adult emergence cages prior to adult eclosion, where they were allowed to emerge over the next 48 h. Newly emerged adult mosquitoes were allowed to mate for at least 48 h and provided with a 10% dextrose solution *ad libitum* via cotton balls. For colony maintenance, female *Ae. aegypti* were fed bovine blood with 0.038% sodium citrate added as an anticoagulant. Blood feeding was performed using parafilm membranes with glass membrane feeders attached to a circulating water bath maintained at 37 °C. Eggs were harvested from 72-96 h post-blood feeding, allowed to embryonate for 48 h and then dried until needed.

### Synthetic Data

**4-((*E*)-((*E*)-1-(*tert*-butyl)-3-methyl-1,3-dihydro-2*H*-imidazol-2-ylidene)triaz-1-en-1-yl)-*N*-(prop-2-yn-1-yl)benzamide (TBD alkyne) (5)**

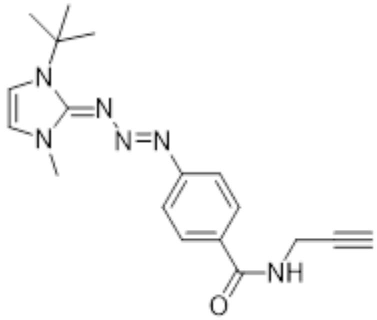

TBD carboxylate[1] (150 mg, 0.49 mmol, 1 eq) was dissolved in 10 mL of $CH_2Cl_2$. Added to this was DIPEA (0.3 mL, 1.7 mmol, 3.5 eq), propargyl amine (150 µL, 2.3 mmol, 4.7 eq) and HATU (186 mg, 0.49 mmol, 1 eq) in order. The reaction was monitored by TLC using a 1 : 1 hexanes : acetone solvent system with neutral alumina TLC plates. The target product is a visibly yellow spot which appears with an Rf ~ 0.3 - 0.5 in this solvent system. The coupling reaction should be completed within 4 h. Next, 10 mL of DI water was added to the reaction mixture which was then stirred vigorously for 20 minutes. A separatory funnel was then used to isolate the product in the $CH_2Cl_2$ layer. The $CH_2Cl_2$ layer is then dried with $MgSO_4$. The crude mixture was filtered, and the filtrate was then adhered onto neutral alumina and purified by running a gradient column (100% hexanes to 1:1 hexanes : acetone). The first yellow spot that comes off the column is not the target product, but the next. An iodine stain should be used to identify all the impurities before re-combining fractions. If product still has left over impurities such as propargyl amine, they can be triturated away using a small amount of $CH_2Cl_2$ followed by a large amount of hexanes. The product is a bright yellow solid (116 mg, 70 %) and stored at RT. Crystals suitable for x-ray diffraction were grown using layering method, with $CH_2Cl_2$ being the solvent and hexanes the precipitant. Vials were left undisturbed for 1 week at room temperature. **$^1$H NMR** (500 MHz, DMSO-$d_6$) δ 8.76 (t, *J* = 5.6 Hz, 1H), 7.80 (d, *J* = 8.5 Hz, 2H), 7.37 (d, *J* = 8.5 Hz, 2H), 7.15 (d, *J* = 2.6 Hz, 1H), 7.00 (d, *J* = 2.5 Hz, 1H), 4.04 (dd, *J* = 5.5, 2.5 Hz, 2H), 3.75 (s, 3H), 3.10 (t, *J* = 2.5 Hz, 1H), 1.62 (s, 9H). **$^{13}$C NMR** (126 MHz, DMSO-$d_6$) δ 165.93, 155.51, 151.42, 129.15, 128.30, 119.71, 118.68, 113.35, 81.63, 72.67, 57.96, 38.49, 28.37, 28.22. **HRMS** (ESI) m/z: [M + H]+ calculated for $C_{18}H_{22}N_6O$ 339.2635; found value 339.1924.

[1] Wijetunge, A. J.; Davis, G. J.; Shadmehr, M.; Townsend, J. A.; Guzmán, L. E.; Marty, M. T.; Jewett, J. C. Copper-Free Click Enabled Triazabutadiene for Bioorthogonal Protein Functionalization, *Bioconjugate Chem.* **2021**, *32*, 254-258.

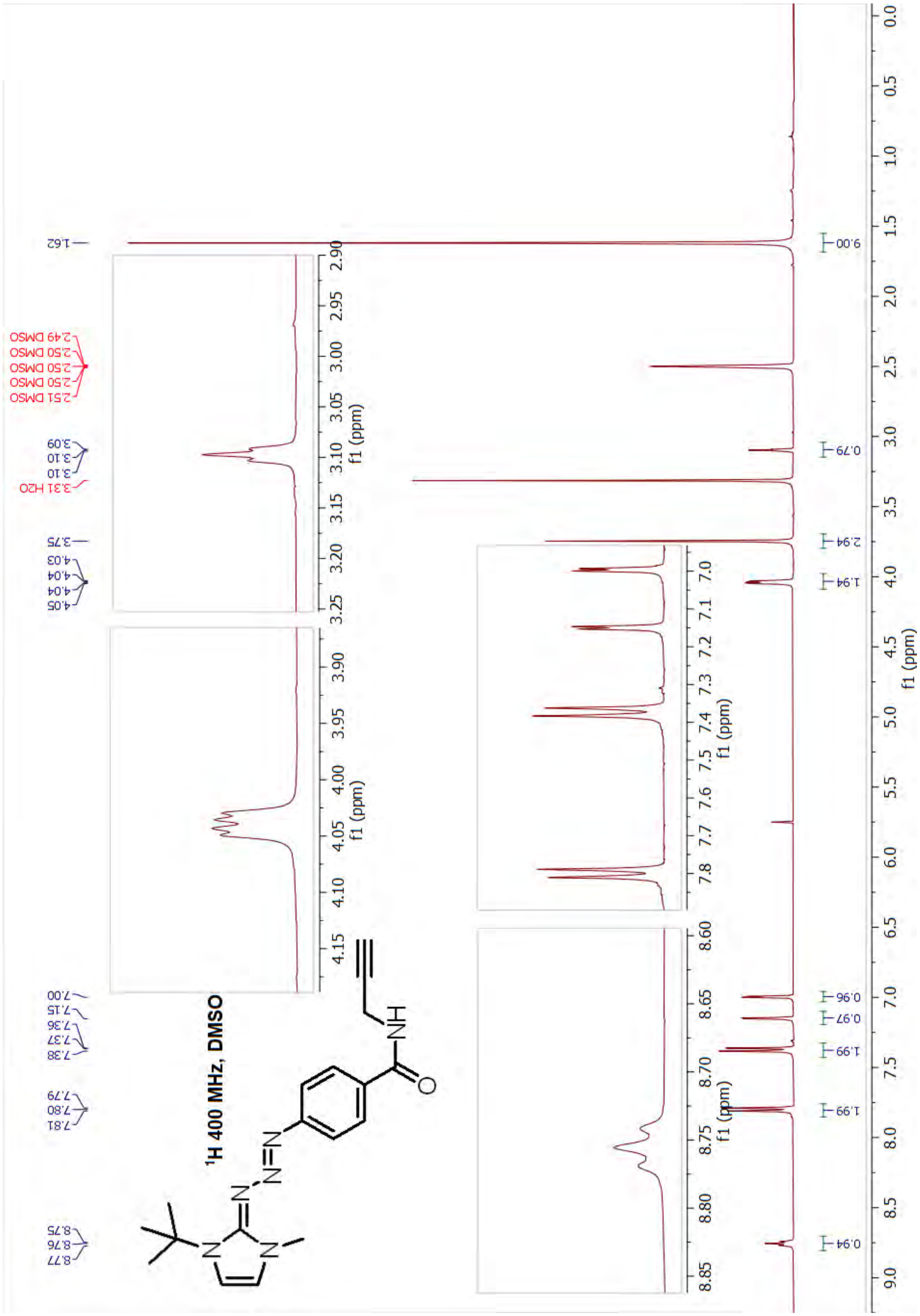

1H 400 MHz, DMSO
1.62
2.49 DMSO
2.50 DMSO
2.50 DMSO
2.50 DMSO
2.51 DMSO
3.09
3.10
3.10
3.31 H2O
3.75
4.03
4.04
4.04
4.05
7.00
7.15
7.36
7.37
7.38
7.79
7.80
7.81
8.75
8.76
8.77
9.00
0.79
2.94
1.94
0.96
0.97
1.99
1.99
0.94
f1 (ppm)

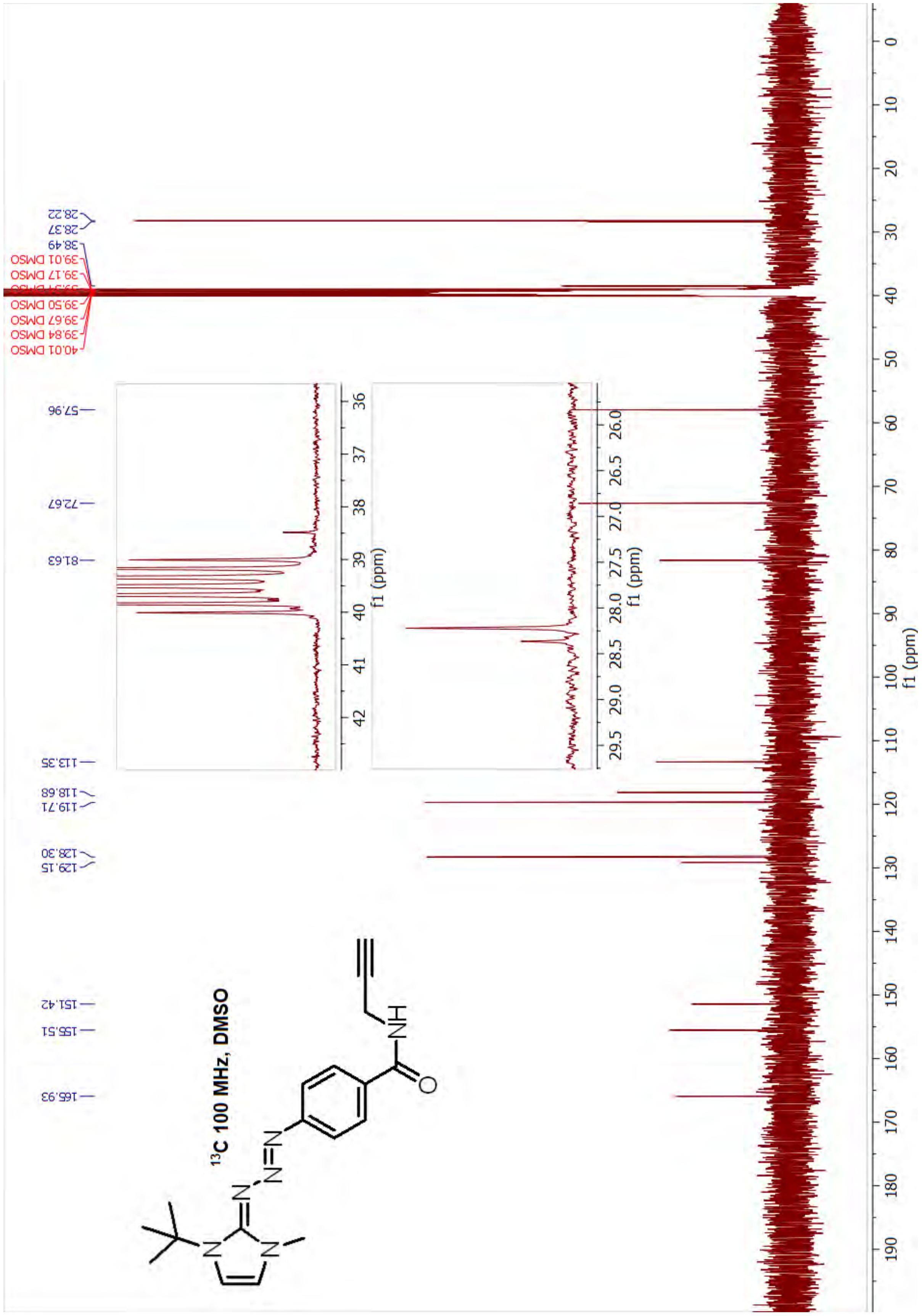
13C 100 MHz, DMSO
28.22
28.37
38.49
39.01 DMSO
39.17 DMSO
39.50 DMSO
39.67 DMSO
39.84 DMSO
40.01 DMSO
57.96
72.67
81.63
113.35
118.68
119.71
128.30
129.15
151.42
155.51
165.93
f1 (ppm)

**(*E*)-3-(*tert*-butyl)-2-(1-(ethoxycarbonyl)-1-(4-(prop-2-yn-1-ylcarbamoyl)phenyl)triaz-2-en-1-ium-3-yl)-1-methyl-1H-imidazol-3-ium chloride (experimental pTBD alkyne) (1)**

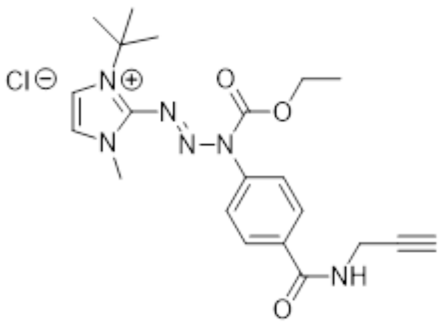

To a flame dried round bottom flask added was anhydrous $MgSO_4$ and 2 mL of anhydrous $CH_2Cl_2$. (the peak at 7.81 ppm can be observed as a doublet with $MgSO_4$ whereas in the previous attempts with molecular sieves it was a singlet). To the same round bottom flask ethyl chloroformate (0.06 mL, 0.6 mmol, 10 eq) was added. After stirring for about 5 min TBD alkyne (**3.1**) (20 mg, 0.06 mmol, 1 eq) was added to the above solution dropwise dissolved in 2 mL of anhydrous $CH_2Cl_2$. Remaining TBD alkyne was washed with another 1 mL of anhydrous $CH_2Cl_2$ and added to the same round bottom flask. The reaction was allowed to stir overnight at RT. The solution was filtered, washed with $CH_2Cl_2$ and the solvent was evaporated. The crude product was obtained as a yellow solid and used without further purification (quantitative). The product was stored at -20 °C. Crystals suitable for x-ray diffraction were grown using the vapor diffusion method, with ethanol being the solvent and diethyl ether the precipitant. Chambers were left undisturbed for about 3-4 weeks at room temperature. **$^1$H NMR** (500 MHz, DMSO-*d*$_6$) δ 9.12 (t, *J* = 5.6 Hz, 1H), 8.07 – 8.01 (m, 2H), 7.81 (d, *J* =0.8 Hz, 2H), 7.58 – 7.52 (m, 2H), 4.43 (q, *J* = 7.1 Hz, 2H), 4.09 (dd, *J* = 5.5, 2.5 Hz, 2H), 3.97 (s, 3H), 3.17 (t, *J* = 2.5, 1H), 1.29 (m, 12H). **$^{13}$C NMR** (126 MHz, DMSO-*d*$_6$) δ 164.86, 151.41, 141.53, 137.87, 135.15, 128.61, 128.54, 123.34, 119.41, 81.14, 73.00, 64.72, 61.96, 38.58, 28.70, 28.59, 14.00. **HRMS** (ESI) m/z: [M]+ calculated for $C_{21}H_{27}N_6O_3^+$ 411.2139; found value 411.2135.

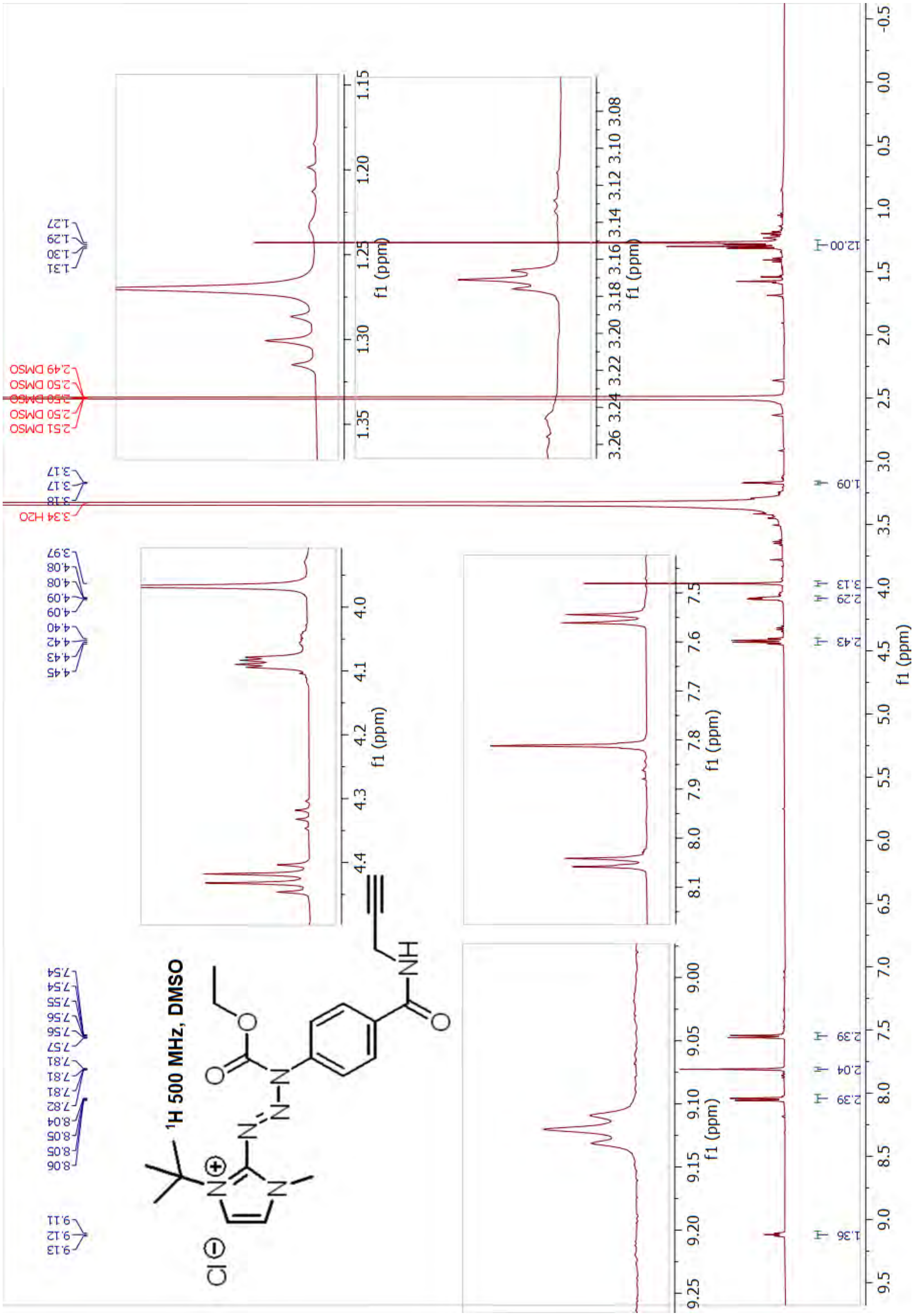

1H 500 MHz, DMSO
f1 (ppm)

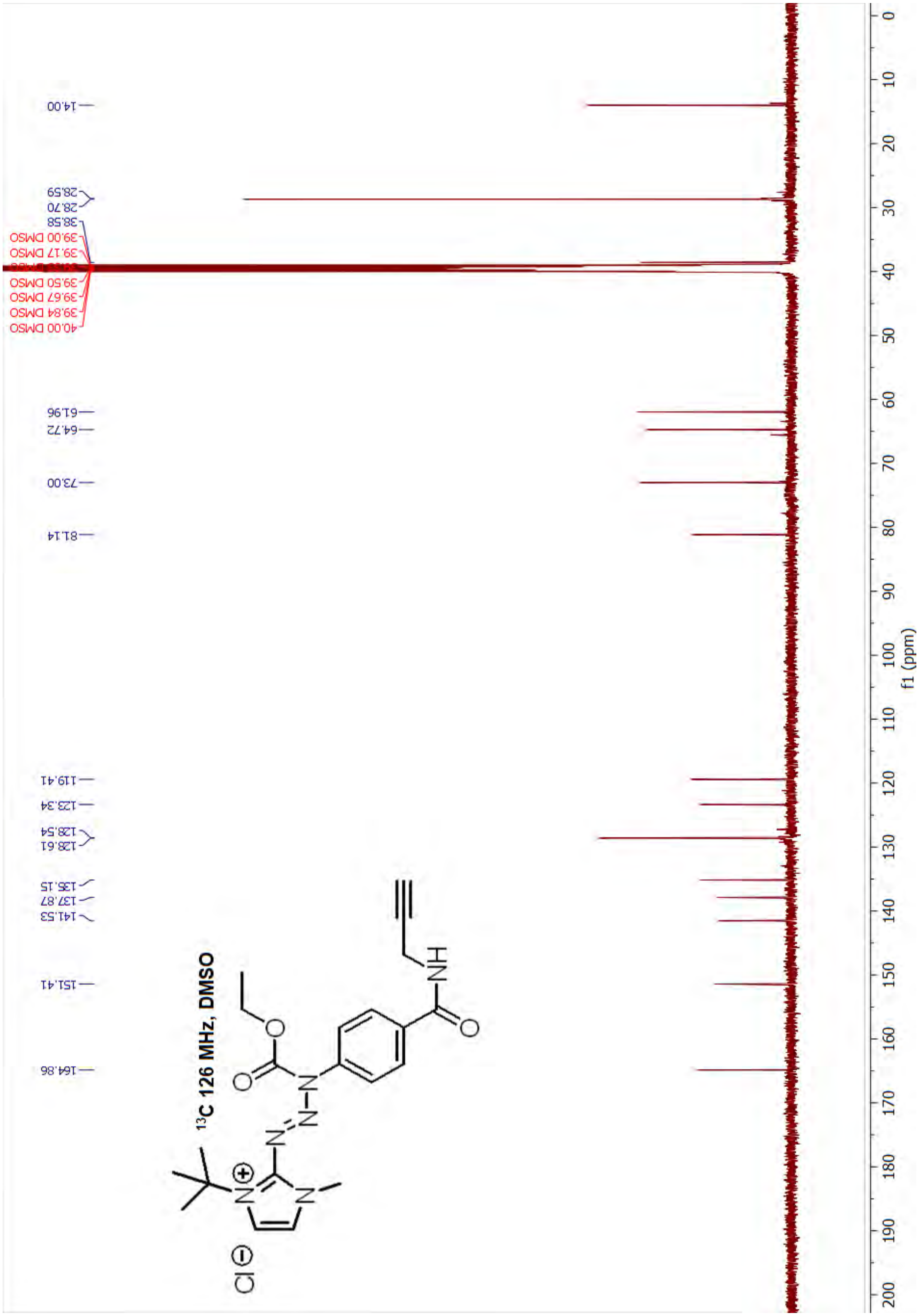

13C 126 MHz, DMSO
14.00
28.59
28.70
38.58
39.00 DMSO
39.17 DMSO
39.50 DMSO
39.67 DMSO
39.84 DMSO
40.00 DMSO
61.96
64.72
73.00
81.14
119.41
123.34
128.54
128.61
135.15
137.87
141.53
151.41
164.86
f1 (ppm)
0 10 20 30 40 50 60 70 80 90 100 110 120 130 140 150 160 170 180 190 200

**(*E*)-3-(tert-butyl)-1-methyl-2-(3-((neopentyloxy)carbonyl)-3-(4-(prop-2-yn-1-ylcarbamoyl)phenyl)triaz-1-en-1-yl)-1*H*-imidazol-3-ium chloride (2)**

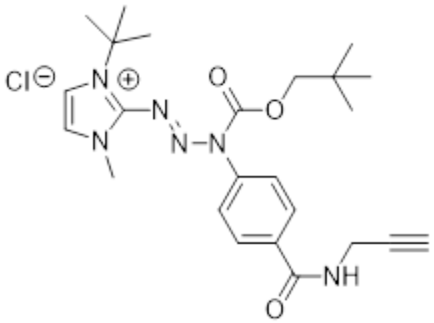

To a flame dried round bottom flask added was anhydrous $MgSO_4$ and 2 mL of anhydrous $CH_2Cl_2$. To the same round bottom flask ethyl chloroformate (225 µL, 1.5 mmol, 10 eq) dissolved in 5 mL anhydrous $CH_2Cl_2$ was added. After stirring for about 5 min TBD alkyne (3.1) (50 mg, 0.15 mmol, 1 eq) was added to the above solution dropwise dissolved in 4 mL of anhydrous $CH_2Cl_2$. Remaining TBD alkyne was washed with another 1 mL of anhydrous $CH_2Cl_2$ and added to the same round bottom flask. The reaction was allowed to stir overnight at RT. The solution was filtered, washed with $CH_2Cl_2$ and the solvent was evaporated. The crude product was obtained as a yellow solid and used without further purification (quantitative). The product was stored at -20 °C. **$^1$H NMR** (500 MHz, DMSO-$d_6$) δ 9.23 (s, 1H), 8.10 (d, $J$ = 8.2 Hz, 2H), 7.87 – 7.81 (m, 2H), 7.57 (d, $J$ = 8.2 Hz, 2H), 4.07 (s, 2H), 4.00 (s, 3H), 3.15 (s, 1H), 3.05 (s, 2H), 1.27 (s, 9H), 0.86 (s, 9H). **$^{13}$C NMR** (126 MHz, DMSO-$d_6$) δ 164.75, 151.33, 141.96, 138.16, 135.79, 128.52, 128.49, 123.32, 119.45, 81.11, 77.05, 72.99, 71.29, 38.55, 32.37, 31.24, 26.25, 25.82. **HRMS** (ESI) m/z: [M]+ calculated for $C_{24}H_{33}N_6O_3^+$ 453.2609 ; found value 453.26091.

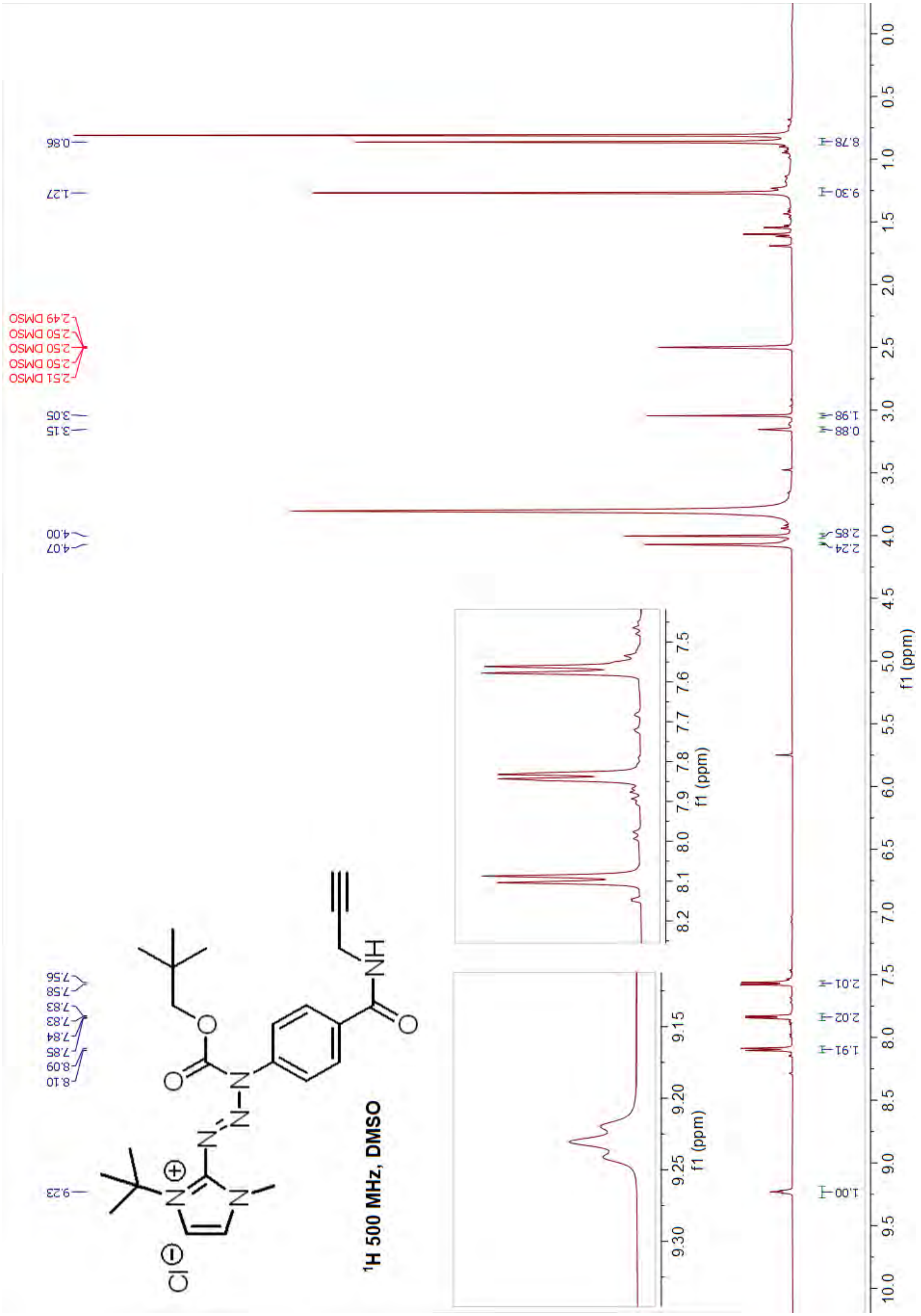
¹H 500 MHz, DMSO
0.86
1.27
2.49 DMSO
2.50 DMSO
2.50 DMSO
2.50 DMSO
2.51 DMSO
3.05
3.15
4.00
4.07
7.56
7.58
7.83
7.83
7.84
7.85
8.09
8.10
9.23
8.78
9.30
1.98
0.88
2.85
2.24
2.01
2.02
1.91
1.00
f1 (ppm)

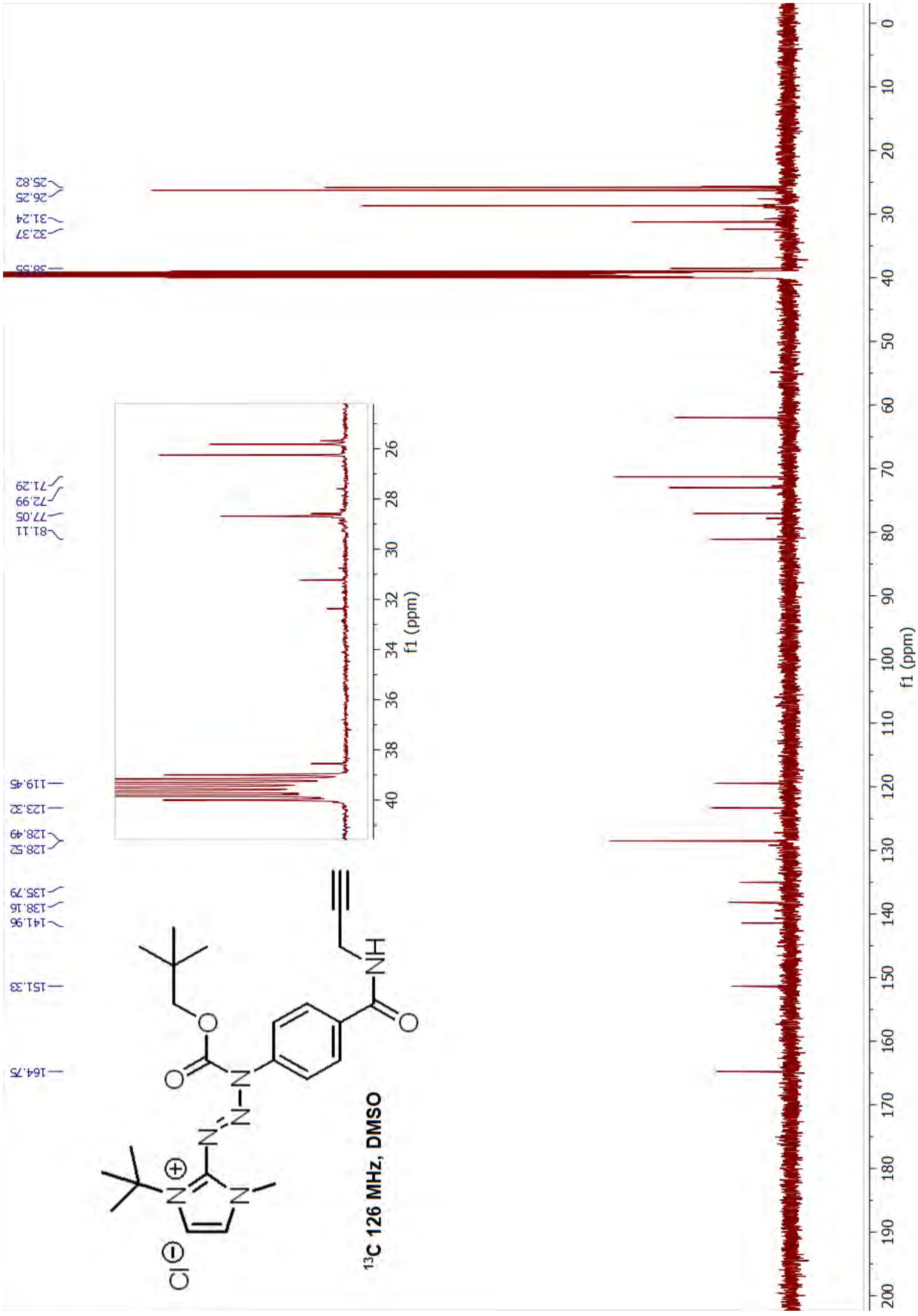

25.82
26.25
31.24
32.37
38.55
71.29
72.99
77.05
81.11
119.45
123.32
128.49
128.52
135.79
138.16
141.96
151.33
164.75
f1 (ppm)
f1 (ppm)
13C 126 MHz, DMSO

**(*E*)-3-(*tert*-butyl)-1-methyl-2-(3-methyl-3-(4-(prop-2-yn-1-ylcarbamoyl)phenyl)triaz-1-en-1-yl)-1H-imidazol-3-ium iodide (control pTBD alkyne) (3)**

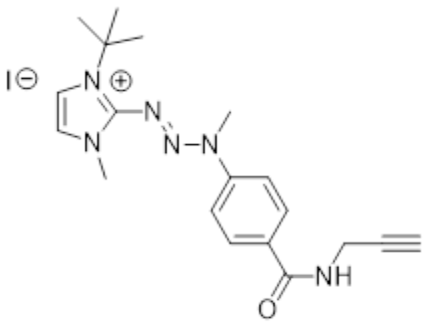

To a flame dried round bottom flask added was 4Å molecular sieves and 2 mL of anhydrous $CH_2Cl_2$. To the same round bottom flask MeI (37 µL, 0.6 mmol, 10 eq) was added. After stirring for about 5 min TBD alkyne (3.1) (20 mg, 0.06 mmol, 1 eq) was added to the above solution dropwise dissolved in 2 mL of anhydrous $CH_2Cl_2$. Remaining TBD alkyne was washed with another 1 mL of anhydrous $CH_2Cl_2$ and added to the same round bottom flask. The reaction was allowed to stir overnight at RT. The solution was filtered, washed with $CH_2Cl_2$ and the solvent was evaporated. The crude product was obtained as a yellow solid and used without further purification (quantitative). The product was stored at -20 °C. Crystals suitable for x-ray diffraction were grown using the vapor diffusion method, with $CH_2Cl_2$/THF being the solvent and hexanes the precipitant. Chambers were left undisturbed for about 3-4 weeks at room temperature. **$^{1}$H NMR** (500 MHz, DMSO-$d_6$) δ 9.04 (t, $J$ = 5.5 Hz, 1H), 8.05 – 7.99 (m, 2H), 7.79 – 7.76 (m, 2H), 7.75 (d, $J$ = 2.4 Hz, 1H), 7.66 (d, $J$ = 2.4 Hz, 1H), 4.08 (dd, $J$ = 5.5, 2.5 Hz, 2H), 3.91 (s, 3H), 3.87 (s, 3H), 3.15 (t, $J$ = 2.5 Hz, 1H), 1.71 (s, 9H). **$^{13}$C NMR** (126 MHz, DMSO) δ 164.85, 145.04, 144.09, 131.75, 128.78, 121.56, 118.80, 117.73, 81.21, 72.98, 61.12, 38.09, 35.55, 28.79, 28.54. **HRMS** (ESI) m/z: [M]+ calculated for $C_{19}H_{25}N_6O^+$ 353.2084; found value 353.2080.

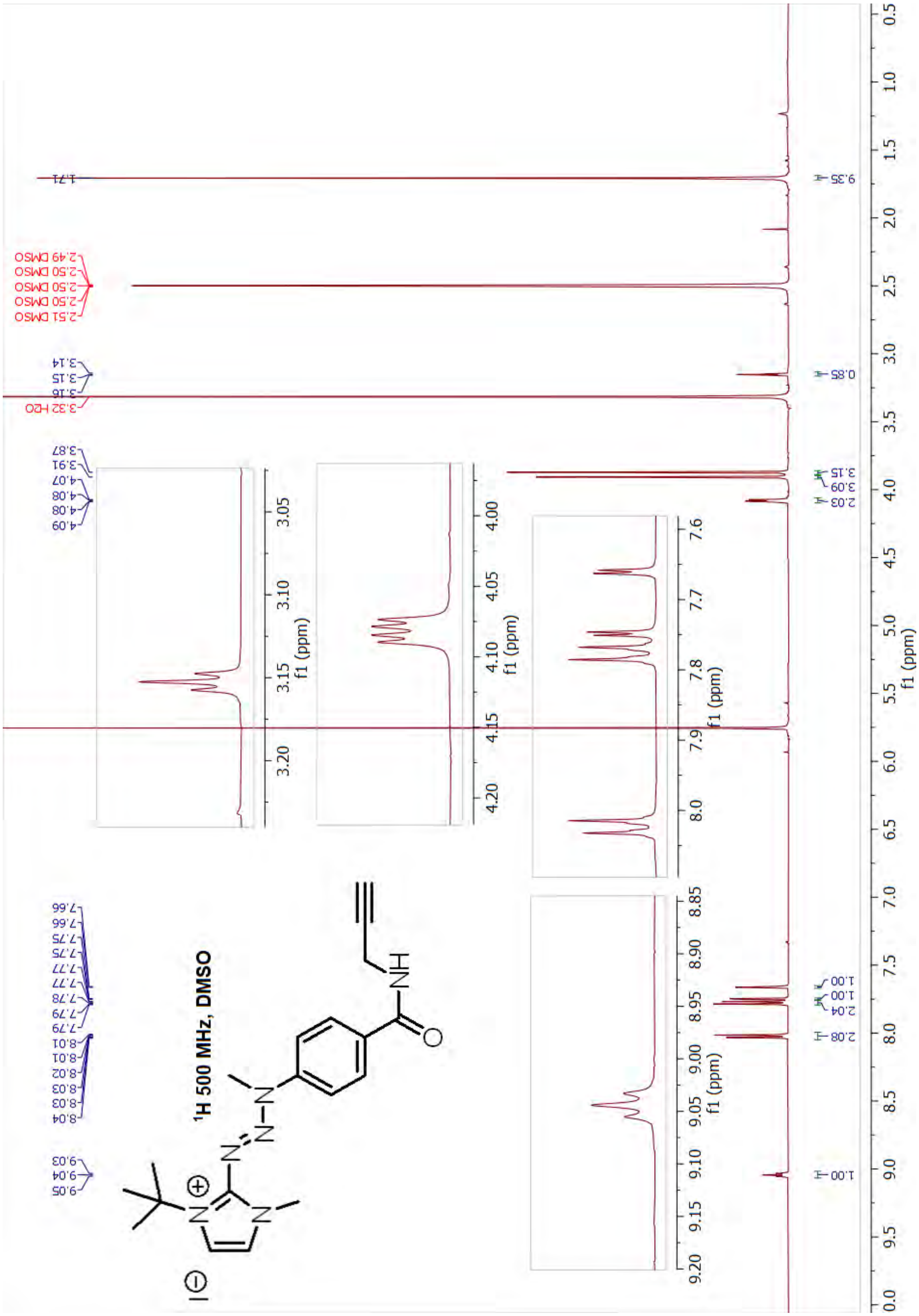

1H 500 MHz, DMSO
f1 (ppm)

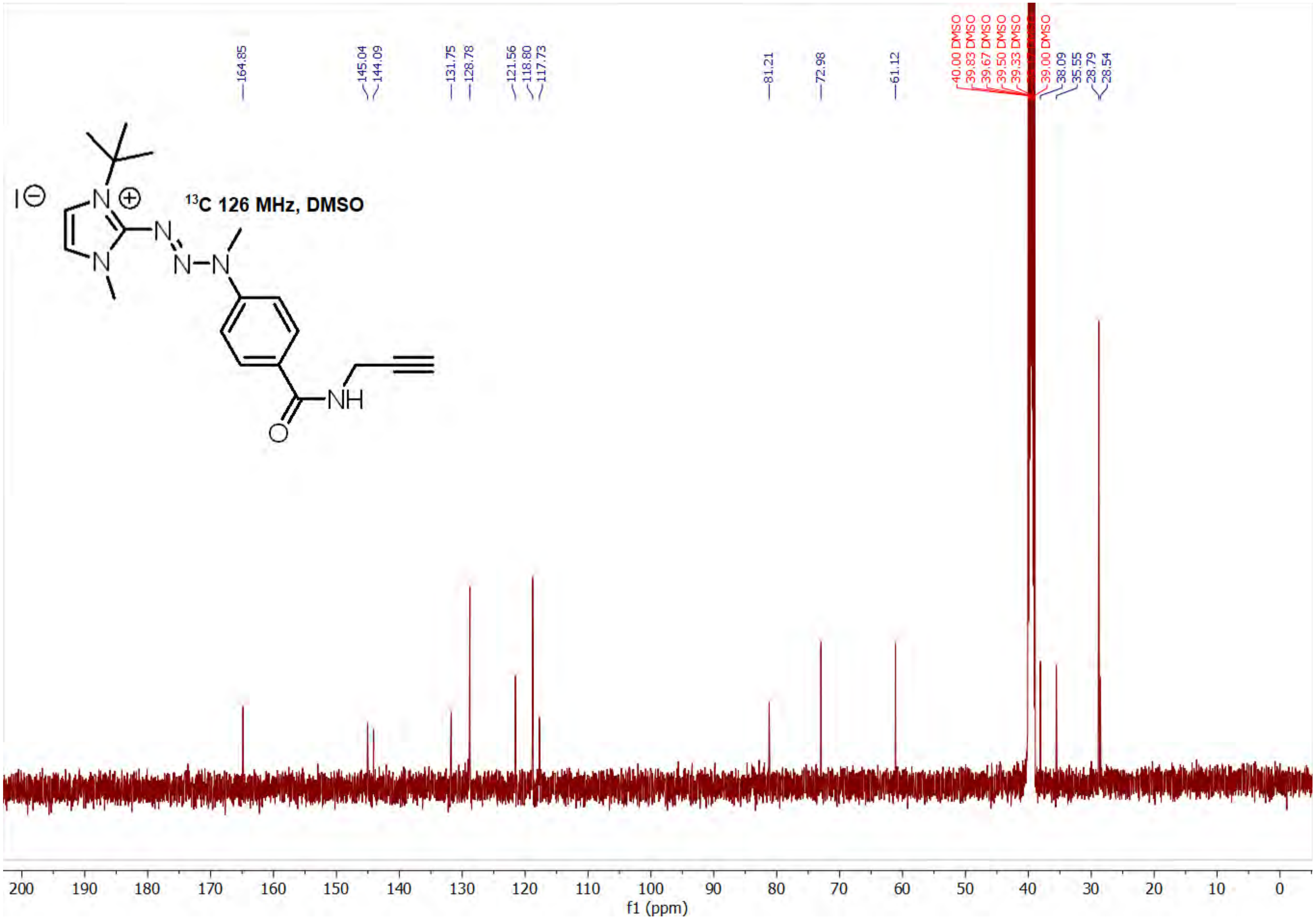

¹³C 126 MHz, DMSO
164.85
145.04
144.09
131.75
128.78
121.56
118.80
117.73
81.21
72.98
61.12
40.00 DMSO
39.83 DMSO
39.67 DMSO
39.50 DMSO
39.33 DMSO
39.00 DMSO
38.09
35.55
28.79
28.54
200 190 180 170 160 150 140 130 120 110 100 90 80 70 60 50 40 30 20 10 0
f1 (ppm)

**Kinetic Evaluation of Triazabutadienes at pH 7 and pH 11**

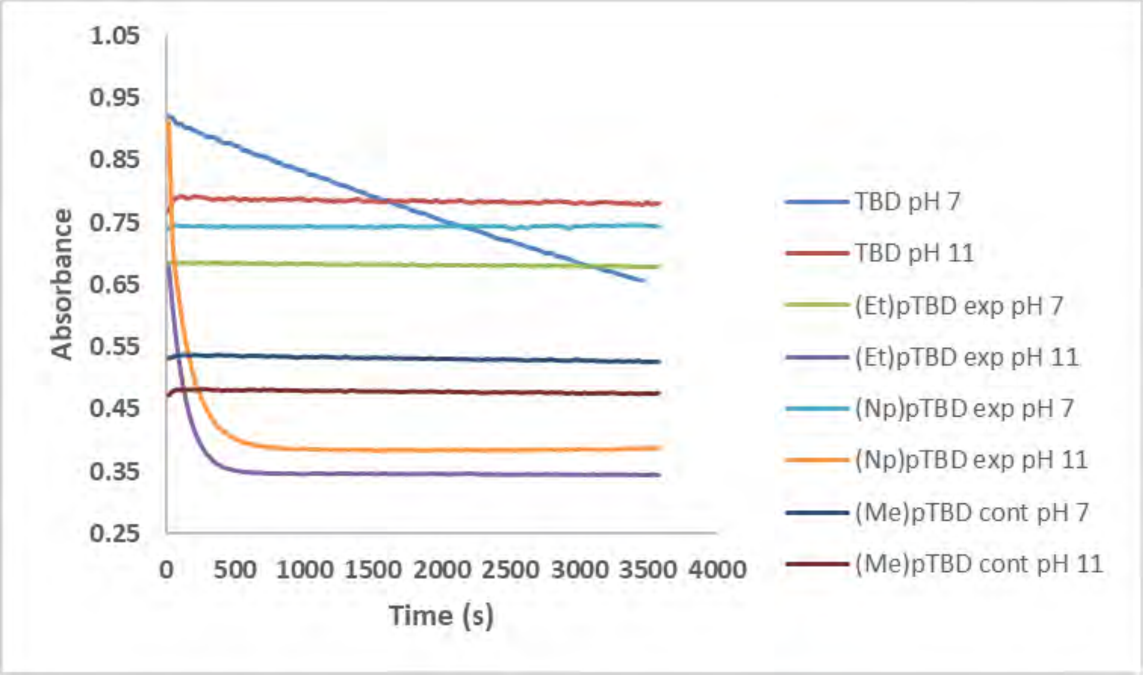

**Figure S1 : The stability studies of (Et)pTBD (1), (Np)pTBD (2), (Me)pTBD (3) and unprotected alkyne-TBD (4) in pH 7 and pH 11 buffers for 1 h.** Absorbance is shown based on the various maximal wavelengths.

### Larval/Biochemical Data

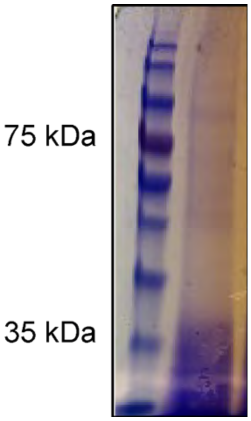

**Figure S2 : Representative Degradation of larval gut proteins upon dissection in general protease inhibitor.** Larvae (20 mosquitoes, 4th instar) were dissected on a glass slide on ice in the presence of 10x of HALT protease inhibitors in PBS. Tissues were then immersed in PBS lyis buffer (1% NP-40, 150 mM NaCl) containing a 10x solution of HALT protease inhibitor. The tissues were then homogenized on ice for 10 minutes and insoluble were removed using centrifugation. Lysate was then incubated at 95 C for 5 minutes in Laemlli sample buffer. Insolubles were removed using centrifugation. The lysate (40 µL) was then loaded onto an SDS-PAGE and run for 45 minutes at 200 V.

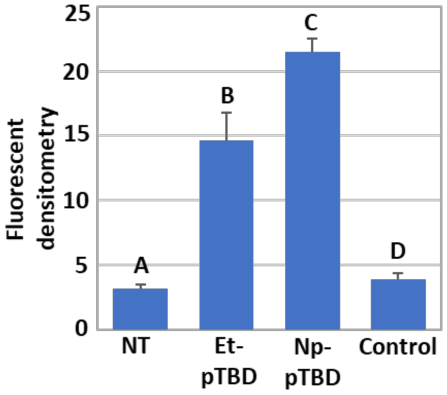

(i)

**Tukey HSD results**

| treatments pair | Tukey HSD Q statistic | Tukey HSD p-value | Tukey HSD inferfence |
|---|---|---|---|
| A vs B | 9.4384 | 0.0010053 | ** p<0.01 |
| A vs C | 15.0513 | 0.0010053 | ** p<0.01 |
| A vs D | 0.6006 | 0.8999947 | insignificant |
| B vs C | 5.6128 | 0.0087219 | ** p<0.01 |
| B vs D | 8.8378 | 0.0010053 | ** p<0.01 |
| C vs D | 14.4507 | 0.0010053 | ** p<0.01 |

(ii)

**Figure S3 : Densitometry analysis of labeled proteins. (i**) Total protein stain using Coomassie brilliant blue to verify equal protein loading between treatments and controls (**ii)** Densitometry analysis of four biological replicates demonstrates a highly significant difference in fluorescent labeling between treatments and controls. Data were analyzed using a one-way ANOVA followed by Tukey HSD post-hoc test. Different letters indicate significant differences at $p<0.01$)

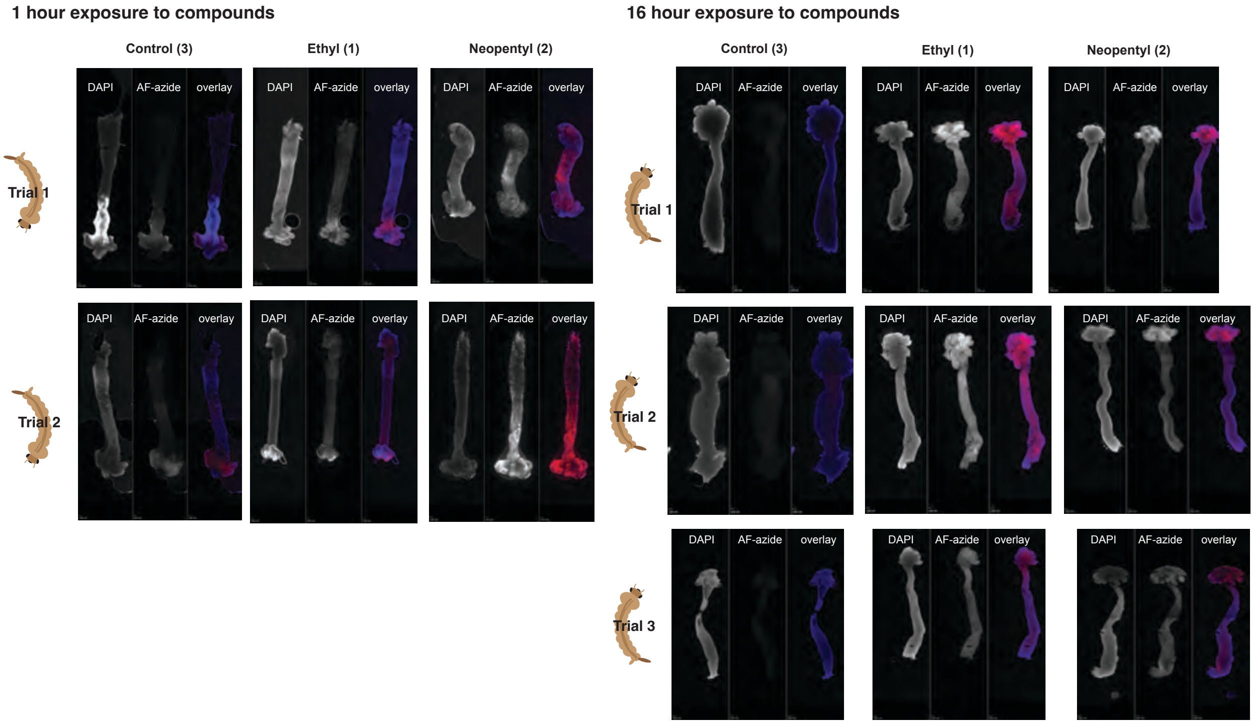

**Figure S4. Larval midgut labeling trials.** All samples were treated and imaged similar to Figure 4 in the manuscript. More variability in labeling in the Az-azide channel is observed in the 1 hour exposure larva than the larva that were treated for 16 hours.

### Sample preparation and performing gel electrophoresis experiments

#### a) Compound treatment and protein extraction for gel electrophoresis

4 petri dishes were obtained. To each petri dish, about 40 *Aedes aegypt* 2nd - 3rd instar larvae were added along with the media. Note that when 4th instar was used upon overnight, they pupated. Each dish was labeled as non-treated (NT), control (C), and experimental (E1 and E2). Next, the media was removed and as soon as the media was removed NT, C, E1 and E2 solutions were added to each of the petri dishes (1 mM, 3 mL). The larvae were allowed to incubate overnight at RT. After overnight incubation, the medium of C and E were replaced with tap water to ensure that the larvae are not exposed to the compounds over the set time. The petri dishes were kept on ice. This makes the larvae immobile and helps in dissections. In the meantime, a 2 mL Dounce homogenizer tube and pestle was prechilled in liquid nitrogen (LN). LN was also poured into the tube directly. Dissections were started with the NT set. To a glass slide kept on an ice block, a 0.1M sodium acetate (pH 3) drop was added. A single larva was placed in the drop and the dissection was performed. Low temperatures and acidic buffer keep the enzymes, such as proteases, inactive and minimizes protein degradation. The gut was separated from the food bolus. 30 guts were dissected per treatment. After each larval dissection, the gut tissue was submerged in LN for a few seconds with the use of a wooden stick appended with a metal probe. Once completely frozen, the tissue was scraped off the stick into the homogenizer containing LN (also submerged in a LN dewer) with the use of another wooden stick. The frozen tissues were ground in the Dounce homogenizer with the prechilled pestle whilst in LN. The Dounce homogenizer was then transferred to the -20 °C along with pestle and tempered for 30 min. It is important to have the setup submerged in LN or -20 °C ice box for transit. This prevents thawing and protein degradation. The tube was then removed from -20 °C freezer. The sample should

look like a nice dry powder at this point. Any thawing will result in protein degradation. Next, the lysis and extraction of the frozen gut tissues were performed as follows. To the homogenizer of powdered gut proteins obtained from -20 °C incubation, solution A was added. Using the pestle, proteins were ground for 1 min at 60 °C (solution A - mixture of 8M urea, 3% SDS and 50 mM tris - 100 µL). Next, solution B was added and the gut proteins were ground for another 4 min at 60 °C with the pestle (solution B – a 1:1 mixture of 0.1 M NaOAc (pH 3) and glycerol - 100 µL). The solution was transferred from the Dounce homogenizer to an Eppendorf tube along with the bubbles and centrifuged at 15000 rpm for 5 min. The supernatant was carefully obtained without disturbing the pellet and aliquoted. Each aliquot was snap frozen in LN. The same above steps were carried out for each compound treated mosquito larval set as well. 30 larval guts were dissected for each treatment and extracted with 200 µL lysis and extraction buffer using the protein extraction protocol stated above (Only needed 30 for dissections but incubated 40 to accommodate any mistakes during the dissection procedure).

**b) Cu Click reaction for gel electrophoresis**

To 10 µL (1.5 guts) aliquots of extracted protein solutions from each treatment, added was the Cu click reaction components according to **Table 1**. The Cu click reaction was carried out for 30 min at RT (total reaction volume upon adding the click reagents = 17.5 µL). The protein samples were mixed with 3 µL of 6X laemmli SDS sample buffer and loaded on a 10 % freshly-made gel (this is a must because fluorescently labeled larval gut lysates behave weird on commercially available gels, especially with the once containing auto fluorescent dyes). The samples were run at 120 V for 1-1.30 h. After fluorescence imaging (546 nm), the gels were stained with CBB G250 overnight, destained and imaged for the presence of proteins.

**Table 1 : Amounts of Cu click reaction components and concentrations used in fluorescence gel electrophoresis studies**

| Reagent | Initial concentration | Volume obtained from stock (µL) | Final concentration |
|---|---|---|---|
| $CuCl_2•2H_2O$ | 15 mM | 1.2 | 1 mM (~3.5 eq) |
| THPTA | 10 mM | 3 | 1.7 mM (~6 eq) |
| AF azide 555 | 5 mM | 1 | 286 µM (~1 eq) |
| Sodium ascorbate | 65.2 mM | 2.3 | 8.6 mM (~30 eq) |

**Sample preparation and performing microscopy experiments**

**a) Compound treatment and protein extraction for microscopy**

A 24 well plate was obtained. The experiment was set in triplicates using 12 wells. To each well *Aedes aegypt* 2$^{nd}$ to 3$^{rd}$ instar larvae were added along with the media. Each well lane was labeled as non-treated (NT), control (C), and experimental (E1 and E2). Next, the media was removed and as soon as the media was removed NT, C, E1 and E2 solutions (1 mM, 500 µL) were added to each of the wells. The larvae were allowed to incubate overnight at RT. After overnight incubation, the medium of C and E were replaced with tap water to ensure that the larvae is not exposed to the compounds over the set time. The well plate was kept on ice. This makes the larvae immobile and helps in dissections. In the meantime, on a separate 24 well plate, 4 wells were labeled as NT, C, and E1, E2. Added to each well was about 500 µL of 4% paraformaldehyde fixing solution (Prepared by mixing 10 mL of 16% paraformaldehyde stock solution with 30 mL of nanopure water. The solution can be stored in -20 °C for several months). Dissections were started with the NT set. To a glass slide kept on an ice block, a 0.1M sodium acetate (pH 3) drop was placed. Larvae was placed on the drop and the dissections were performed. Low temperatures and acidic buffer are essential to keep the enzymes, such as proteases, inactive which reduces protein degradation. The gut was separated from the food bolus. As soon as the guts were dissected, they were immersed in the paraformaldehyde solution in the well plate that was kept on ice. Do not use more than 6 guts in one well (the experiments were done in 24 well plates and having too many larvae per well can damage the guts during washing steps). Likewise, the C and E samples were also subjected to dissections as stated and immersed in 4% paraformaldehyde. The fixation was done at 4 °C overnight. The following day, washing steps and tissue permeabilization was carried out according to the following order, a) Paraformaldehyde was removed from each well and the guts were washed with 500 µL of 1X PBS (3x10 min) at RT. b)The guts were treated with 500 µL of 250 mM glycine solution for 30 min at RT to quench any remaining paraformaldehyde. c)The guts were further washed with 500 µL of 1X PBS (3x5 min) at RT. d)The guts were resuspended in 500 µL of 0.1% Triton-X in 1X PBS for 1h at RT. Triton-X is a permeabilizer and allows creation of the pores in the fixed tissue to allow the penetration of small molecule fluorophores while retaining the fixed 3D architecture. e)The gut tissues were resuspended in 500 µL 1X PBS until the rest of the steps are performed.

**b) Cu Click reaction for microscopy**

Cu click chemistry protocol for staining fixed/ permeabilized cells was used with slight modifications. $CuSO_4$ and sodium ascorbate were prepared fresh each time. *https://clickchemistrytools.com/product/afdye-555-azide-plus/ (Nano pure or DI water was used for solution preparation).*The protocol was first accessed on 03/25/2021.

**Table 2 : Amounts of Cu click reaction components and concentrations used for fluorescence imaging of the mosquito larval guts**

| Reagent | Stock concentration | Volume obtained from stock (µL) |
|---|---|---|
| $CuSO_4 \cdot 5H_2O$ | 50 mM | 5 |
| THPTA | 10 mM | 0.2 |
| AF 555 azide plus | 5 mM | 0.2 |
| Sodium ascorbate | 100 mM | 50 |
| 1X PBS | | 444.6 |
| | | Total vol = 500 |

1X PBS was removed from all the wells (NT, C, E1, E2) and the click cocktail was added. 444.6 µL of 1X PBS was added to each well and the rest of the above solutions were added in the following order. ($CuSO_4$, THPTA, AF 555 azide fluorophore, sodium ascorbate so that the total volume becomes 500 µL). The Cu click reaction was performed for 30 min. The guts were washed with 500 µL of 1X PBS (2x10 min) and (1x24 h) to get rid of the excess fluorophore. The guts were carefully placed on a glass slide side by side on a drop of the antifade mounting media containing DAPI. Then a coverslip was laid on top of the specimen without creating any bubbles. The slide was kept in dark for 24 h for complete staining and then imaged.

**Analysis of the components in the lysis and extraction buffer and their effect on the Cu click reaction**

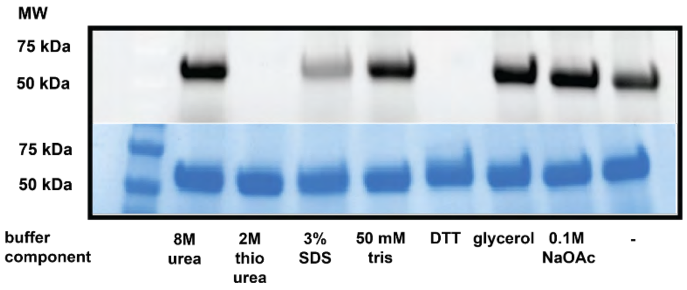

**Figure S3 :: Effect of the lysis and extraction buffer on the Cu click reaction. Thiourea and DTT completely inhibits the Cu click while SDS also has a certain effect for reaction inhibition.**

To 352 µL of 0.1M PBS 7, 40 µL of 250 µM of BSA prepared in 0.1M PBS 7 was added. To this added was 8 µL of 20 mM unprotected TBD alkyne (compound 1) prepared in DMSO. The labeling reaction was allowed to occur for 1h at RT on the shaker. Next, 100 µL of the above solution was taken and 4 µL of 0.5M resorcinol prepared in DI water was added. The reaction was allowed to occur for 1h at RT on the shaker. This step was done to quench any excess unprotected TBD alkyne thus ensuring that there is no more labeling. Next, to 10 µL aliquots from the above step, added was 1 µL of each lysis and extraction buffer component (8M urea, 2M thiourea, 3% SDS, 1 mg DTT in 0.1M PBS, 0.1M NaOAc, glycerol) separately. A control reaction was done where 1 µL of 0.1M PBS 7 was mixed with 10 µL of resorcinol treated protein above. To each of the above reactions Cu click reagents were added according to the **Table 1**. The click reaction was allowed to occur for 30 min at RT (total reaction volume = 18.5 µL). Then each test and control samples were mixed with 3 µL of 6X laemmli SDS sample buffer and loaded on a gel. The samples were run at 120 V for 1h.

**Supporting Video 1**: Larval guts, shown in **Figure 4A** where imaging with spinning disc confocal microscopy to provide Z-stacking cross section videos as described in the **General Information** section (above).

**Supporting Video 2**: Larval guts, shown in **Figure 4B** where imaging with spinning disc confocal microscopy to provide Z-stacking cross section videos as described in the **General Information** section (above).
